# Single-Cell Data Analysis Using MMD Variational Autoencoder for a More Informative Latent Representation

**DOI:** 10.1101/613414

**Authors:** Chao Zhang

## Abstract

Variational Autoencoder (VAE) is a generative model from the computer vision community; it learns a latent representation of images and generates new images in an unsupervised way. Recently, Vanilla VAE has been applied to single-cell data analysis, in the hope of harnessing the representation power of latent space to evade the “curse of dimensionality” of the original dataset. However, Vanilla VAE is suffering from the issue of less informative latent space, which raises a question concerning the reliability of Vanilla VAE latent space in representing the high-dimensional single-cell datasets. Therefore I set up such a study to examine this issue from the multiple perspectives.

This paper confirms the issue of Vanilla VAE by comparing it with MMD-VAE, a variant of VAE which has claimed to have overcome this issue based on image data, across a series of single-cell RNAseq and mass cytometry datasets. The result indicates that MMD-VAE is superior to Vanilla VAE in retaining the information not only in the latent space but also the reconstruction space, which suggests that MMD-VAE be a better option for single-cell data analysis than Vanilla VAE.

## 1 Introduction

In recent years, deep learning has achieved much success in computer vision, speech recognition, natural language processing, audio recognition and so on. With this influence, deep learning has begun to percolate into many computational areas in computer science like probabilistic graphical models. Variational Autoencoder (VAE)[1] is such a product of deep learning and probabilistic graphical models. VAE has a deep neuron structure similar to autoencoder (see Figure 1) on the one hand and is a probabilistic model in terms of original space (data *x*) and latent space (latent variables *z*) on the other hand[2] (Supplementary Figure S1).

**Figure 1:**
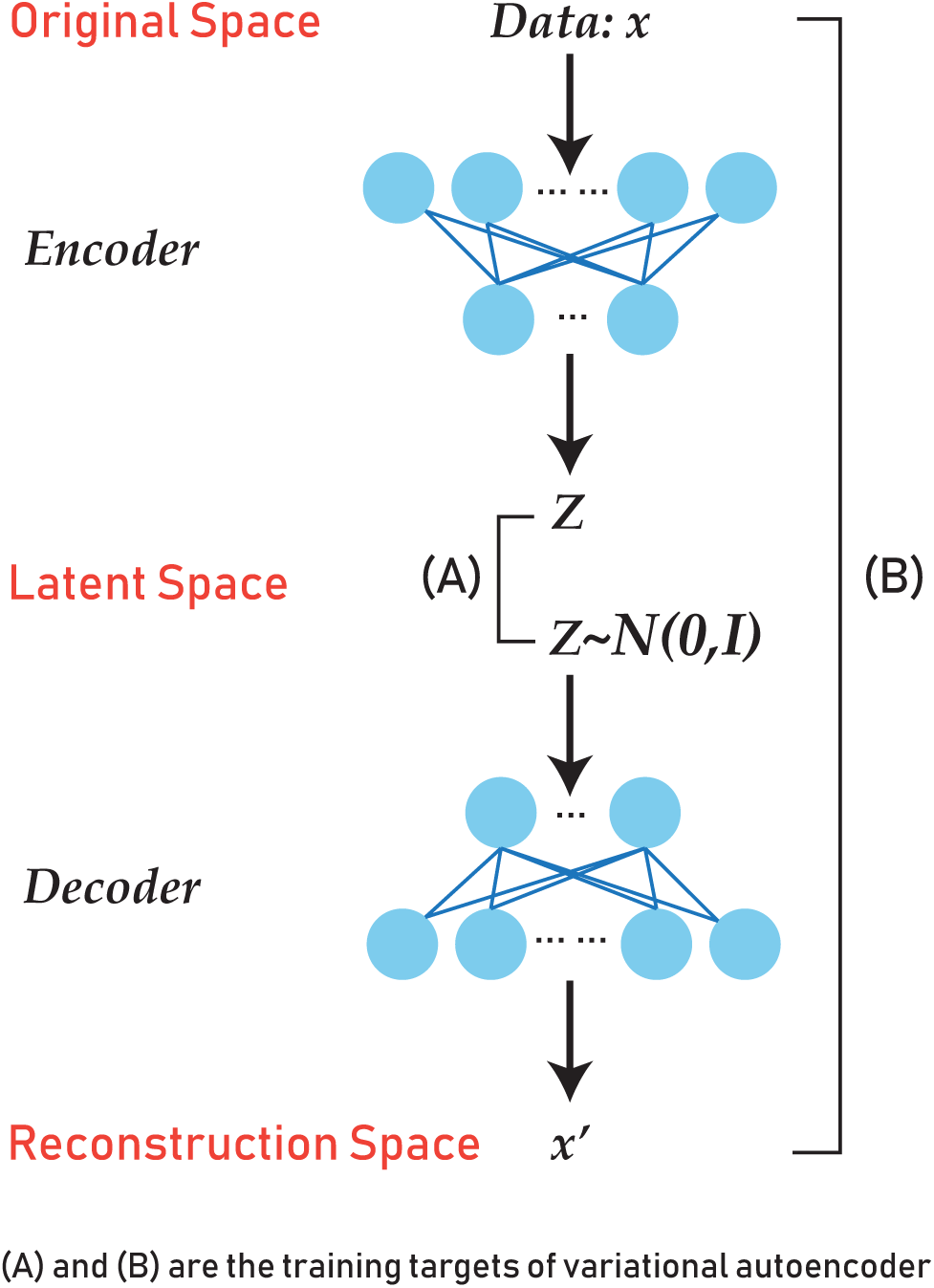
An intuitive representation of variational autoencoder. (A) and (B) comprises the training target: (A), *q*_*ϕ*_(*z*) should be as similar as possible to *p*(*z*), the simple distribution, say, a normal distribution; (B), the original space *P*(*X*) and reconstruction space *P*(*X*′) should be the same.

The flexibility of VAE architecture makes it possible to use VAE in different ways. The most obvious one is the generative purpose. For example, VAE can learn a latent space *Z* from the existing images *X* and generate new images *x*′ that make sense to humans by randomly sampling the learned latent space *Z*. Besides, the latent space and reconstruction space is also worth exploration under proper usage scenario. This has sparked some interest from other research community in applying VAE.

In recent years, the rapid development of biology experiment techniques like single-cell RNAseq[4] and mass cytometry[3] has led to a massive amount of high-throughput molecular data on a single cell level. Single-cell RNAseq has been able to capture the RNA expressions on a single cell level, which provides new possibilities in understanding the biological activities. Currently, the analysis of single-cell RNAseq data still revolves around the classifying and subsetting cell population in order to depict the cell heterogeneity[6] or constitution[7]. Mass cytometry is a mass spectrometry technique to determine the properties of cells, where antibodies are used as probes to label cellular proteins. Manual gating has been extensively used to determine new cell types from cytometry dataset, but the high dimensions make this operation less practical, especially when to discover new rare cell types[5]. The analysis of multiple-cell high-dimensional datasets has challenged the capacity of the classical unsupervised learning methods, and therefore requires the development of new computational methods to deal with the complexity.

As an unsupervised learning method, the usage of VAE can be viewed from another perspective; that is, VAE learns a latent space *Z* that can represent the original space *X*. In particular, if *x* is high-dimensional and *z* is a low-dimensional, VAE seems to border on dimension reduction and feature extraction, the traditional unsupervised learning strategy. This potential of VAE concurs with the interests of the bioinformatics community regarding reducing the dimensions of the multi-sample high-dimensional datasets for analysis. As a consequence, there have been a few explorative attempts[8][9][10][11] at applying VAE to single-cell datasets, hoping to harness of the low-dimensional representation of the latent space to evade the “curve of dimension” of high-throughput data.

Vanilla VAE is usually considered as the first choice when applying it to other domains like bioinformatics because it has been studied more thoroughly than its variants. However, previous computer vision research in VAE method has shown that Vanilla VAE suffers from the issue of less informative latent space[12][13][14], which means the latent space might not be so meaningfully representative of the original space, potentially undermining the reliability of using VAE as a dimension reduction method. Meanwhile, MMD-VAE, using Maximum Mean Discrepancy (MMD) instead of Kullback–Leibler divergence as part of the training target, has overcome this issue[12], looking promising as an alternative to Vanilla VAE in bioinformatics and computational biology research. Previous research was done based on image data, which can be validated directly by mesh eyes; however, biological data are very different, and the validation strategy for image data does not apply to biological data; therefore something has to be done to bridge the gap.

In this paper, I carried out an experimental study to examine the issue of less informative latent space issue using biological datasets as opposed to image datasets, in order to bridge the gap between the instinct of exploiting VAE latent space on the image datasets and that on the biological datasets. Briefly, the study compared Vanilla VAE and MMD-VAE based on the single-cell datasets of RNAseq and mass cytometry by adopting a validation scheme which involved separate neural network classifiers, confirmed the issue on Vanilla VAE and further demonstrates the superior representativeness of latent space of MMD-VAE over Vanilla VAE.

## Results

### Validation scheme

To compare and evaluate Vanilla VAE and MMD-VAE over single cell data, I adopted a validation scheme as Figure 2 shows. For each dataset, I trained a neural network classifer separately over each VAE space with the same labels. The accuracy over the test data in VAE latent and reconstruction space was used as a measurement of how much information is retained, compared to the accuracy over the test data in VAE original space.

**Figure 2:**
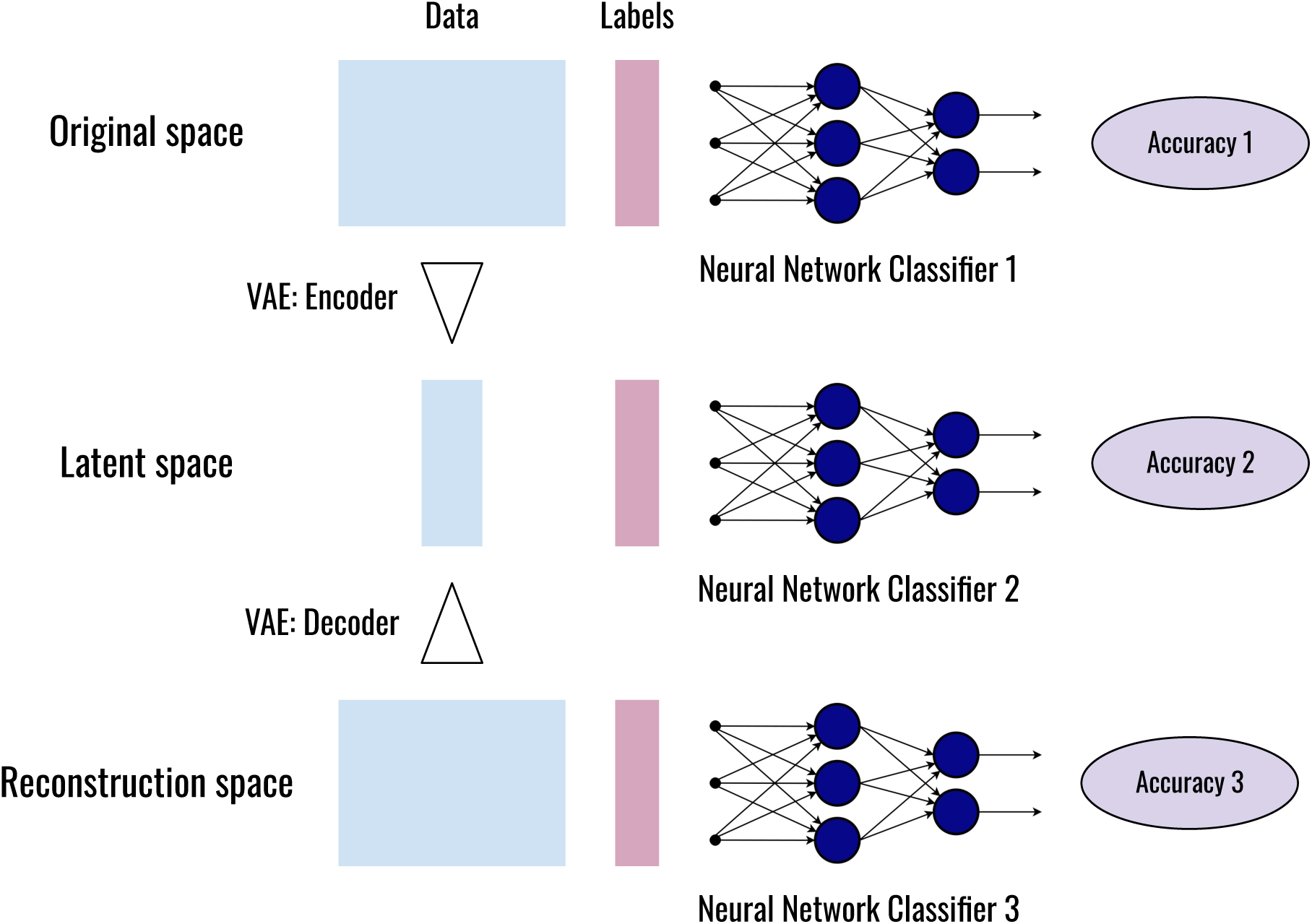
The validation scheme to compare Vanilla VAE and MMD-VAE over single cell datasets. For each dataset, 3 neural classifiers were applied to original space, latent space and reconstruction space separately, ending up with 3 accuracies. The accuracy over the test data in VAE latent and reconstruction space was used as a measurement of how much information is retained in each space, compared to the accuracy over the test data in VAE original space.

### Classifier performance

The neural network classifier was applied to the original space of four datasets separately, RBN, AML and PAN have achieved a high accuracy (Supplementary Table S1), while SN has achieved a moderate accuracy, which could be due to the insufficient samples (only 622 cells) and the different unsupervised methods used by the experts in the original papers to classify the datasets; the moderate accuracy here may reflect the disagreement between the current classifier and the past unsupervised method. The accuries I got here serves as a baseline to evaluate the retained information in latent and reconstruction space.

With regards to the classifier accuracy over the latent space of VAE, MMD-VAE outperforms Vanilla VAE across all the single-cell datasets, as Table 1 shows; incidentally, with regards to the classifier accuracy over the reconstruction space of VAE, MMD-VAE still outperforms Vanilla VAE across all the single-cell datasets (Table 2).

**Table 1:** The accuracy of neural network classifier over the latent space of VAE (the average accuracy+/-standard deviation).

| Dataset | Metrics | vanilla VAE | MMD-VAE |
| --- | --- | --- | --- |
| SN | F2 | 0.364+/-0.064 | 0.576+/-0.041 |
|  | MCC | 0.415+/-0.045 | 0.658+/-0.027 |
| RBN | F2 | 0.901+/-0.032 | 0.964+/-0.007 |
|  | MCC | 0.924+/-0.020 | 0.968+/-0.006 |
| AML | F2 | 0.869+/-0.026 | 0.896+/-0.021 |
|  | MCC | 0.891+/-0.030 | 0.917+/-0.009 |
| PAN | F2 | 0.846+/-0.033 | 0.925+/-0.005 |
|  | MCC | 0.870+/-0.029 | 0.933+/-0.005 |

**Table 2:** The accuracy of neural network classifier over the reconstruction space of VAE (the average accuracy+/-standard deviation).

| Dataset | Metrics | vanilla VAE | MMD-VAE |
| --- | --- | --- | --- |
| SN | F2 | 0.454+/-0.050 | 0.752+/-0.024 |
|  | MCC | 0.460+/-0.047 | 0.743+/-0.024 |
| RBN | F2 | 0.928+/-0.027 | 0.974+/-0.003 |
|  | MCC | 0.941+/-0.019 | 0.977+/-0.003 |
| AML | F2 | 0.873+/-0.040 | 0.905+/-0.015 |
|  | MCC | 0.893+/-0.035 | 0.922+/-0.008 |
| PAN | F2 | 0.857+/-0.038 | 0.926+/-0.007 |
|  | MCC | 0.875+/-0.032 | 0.933+/-0.006 |

### Performance of VAE

The executing time of MMD-VAE is a little longer than Vanilla VAE (Figure 3), due to the relatively expensive calculation of MMD, if not the problem of code efficiency. In practice, across all the four datasets, the difference of the running time between two VAE architectures lies between 10 and 30 seconds. The reconstruction loss of MMD-VAE is generally smaller than that of Vanilla VAE in the experiment (Figure 4), which may correspond to the distinct classifier accuracies over the reconstruction space.

**Figure 3:**
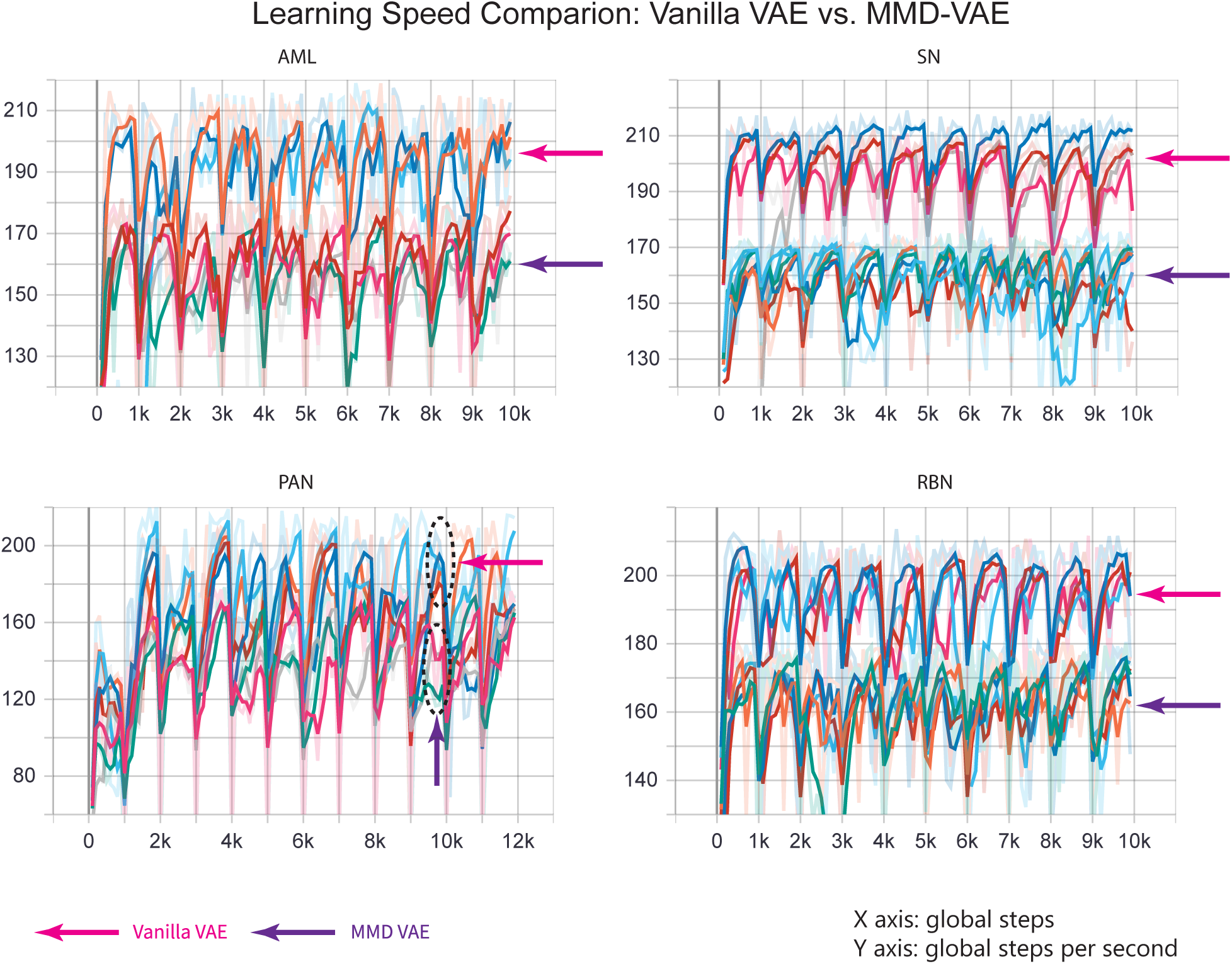
The training speed comparison between Vanilla VAE and MMD-VAE across four single-cell datasets. Each of the colourful lines represents a VAE run. The only difference between these two VAE architectures is the divergence. the calculation of MMD is more expensive, but thanks to the kernel embedding trick, the time it takes in practice is only a little longer than KL divergence. In the experiment, across all the four datasets, the difference of the running time between two VAE architectures lies between 10 and 30 seconds.

**Figure 4:**
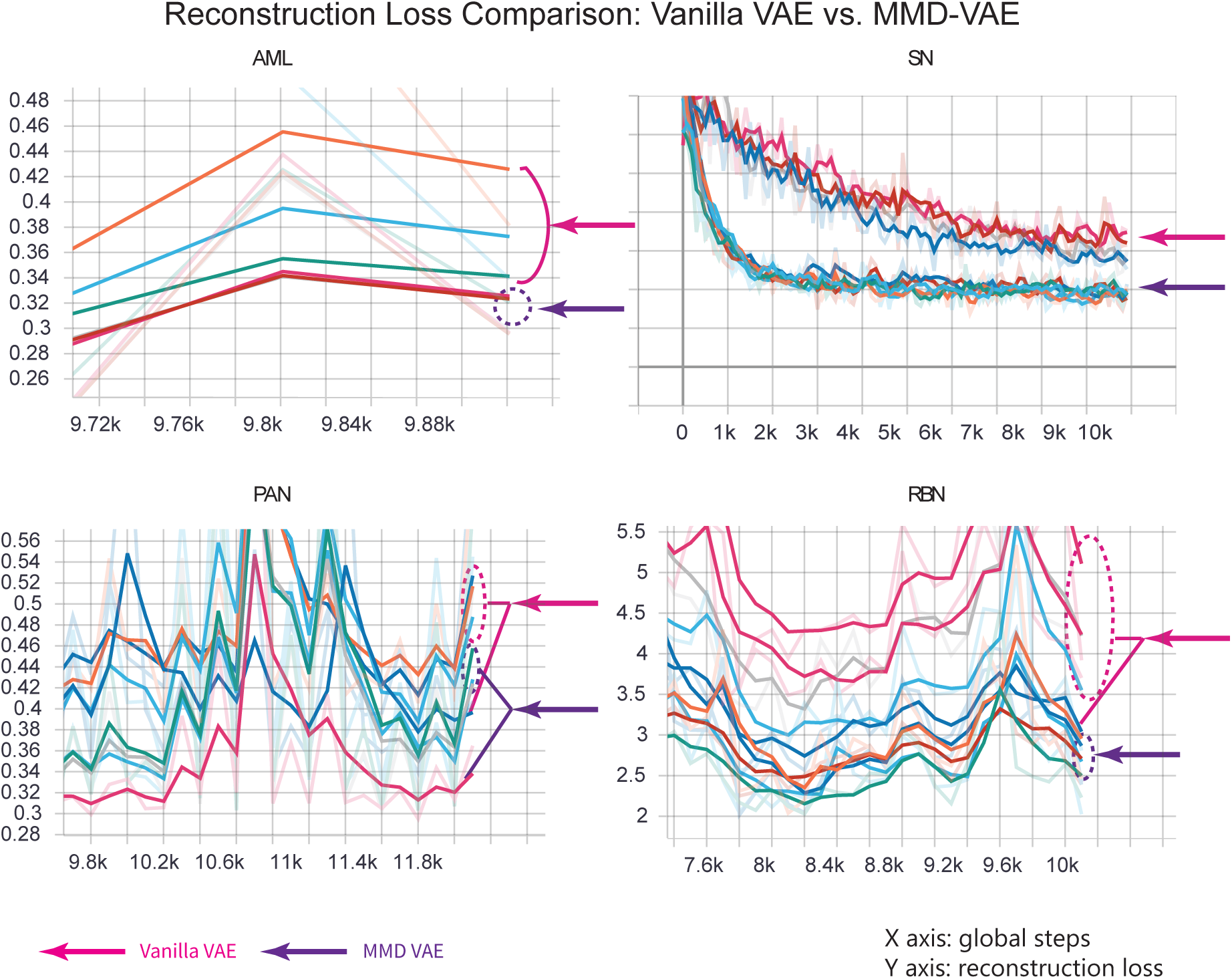
The reconstruction loss comparison between Vanilla VAE and MMD-VAE across four single-cell datasets. Each of the colourful lines represents a VAE run. MMD-VAE in general has a lower reconstruction loss than Vanilla, which may correspond to the distinct classifier accuracies over the reconstruction space.

Besides, in the current experiment, MMD-VAE has less unlucky runs (1 unlucky run out of 23 runs across four datasets), those runs with worse performance (in terms of the accuracy of neural network classifier), than Vanilla VAE (6 unlucky runs out of 28 runs across four datasets). This observation may indicate that MMD-VAE is less subject to the stochastic property of architecture than Vanilla VAE; that is, MMD-VAE is more stable in getting a meaningful latent representation of the data.

## Discussion

Compared to the classifier accuracy achieved in the original datasets, in the reconstruction space and latent space of (Vanilla/MMD) VAE, the accuracy is lowered a little bit. This difference may be attributed to the nature of VAE architecture, which is pursuing an approximation, not exactness. This difference may also respond to the observation when VAE is applied to image data and that the newly generated image is still blurred, which probably means lost of information is inevitable. For those numbers in the biological datasets, the meaning usually does not come from the exactness of numbers, but the comparison between numbers; therefore the lowered accuracy may not hurt a thing when it comes to interpreting the data.

In this study, one challenge is to measure how much information is retained in the latent space of VAE, which is most relevant to the bioinformatics research. Considering the widespread practice of the current biological research, I assume that the information is “materialized” by the labels created by human experts in the given datasets. This information should be kept in latent space and reconstruction space. To measure how much information is retained, I chose to use a neural network classifier to classify the latent/reconstruction space using the labels. since it only works on the true data (Figure 2), not on the permuted data as my separate permutation experiment shows.

The comparison of two VAE architectures concerning how much information is retained is tricky because many subjective factors need to be taken into consideration. The (hyper)parameters of VAE architectures and three neural network classifiers (Figure 2) per each dataset need to be optimized; the stochastic nature of neural networks and the complexity of datasets need to be considered. Therefore it is essential to have a proper mental model to guide the design of the experiment. I used a strategy similar to the philosophy behind statistical hypothesis testing in this study: I had assumed that MMD-VAE should be no worse than Vanilla VAE, so I optimized only the parameters of Vanilla VAE and its corresponding neural network classifier for each dataset, the result only reflects the efforts I had made to optimize, not the best possible result, which I couldn’t guarantee to achieve. I reused these parameters for MMD-VAE and its corresponding three neural network classifiers accordingly; then I obtained the result, that’s the accuracies in terms of the two metrics. As long as the result shows that MMD-VAE outperforms Vanilla VAE, it corroborates the assumption and this experiment strategy works; if not, this would only mean that MMD-VAE is no better than Vanilla VAE under this mental framework.

The result indicates the advantage of MMD-VAE over Vanilla VAE in retaining information of interest; MMD-VAE tends to run a little slower than Vanilla, due to the calculation of MMD. In practice, this may not be a big issue compared to the advantage of MMD-VAE over Vanilla VAE. In my experiment, MMD-VAE revealed a more stable performance than Vanilla VAE. For the reconstruction space, which is less of a focus in this paper, MMD-VAE also outperforms Vanilla VAE, well corresponding to the computer vision research where the generated image from MMD-VAE seems to be sharper than Vanilla VAE[15]. It may also correspond to the lower reconstruction loss of MMD-VAE (Figure 4). The reconstruction space seems to be worth further exploration, though the usage scenario is not explicit at first sight.

Admittedly, this research only covers the usage scenario that VAE is directly applied to datasets without further customization and change in its architecture (which would require extra explorations) in order to keep a comprehensive comparison plausible, but I hope this comparison between MMD-VAE and Vanilla VAE can still give some intuition to people who intend to apply VAE to biological datasets in a more sophisticated way.

In summary, the issue of less informative latent space concerning Vanilla VAE probably limits the exploitation of its latent space to interpret the high-dimensional single-cell datasets; meanwhile, MMD-VAE does not have this issue. From these comparisons, I may draw the following conclusion: MMD-VAE is a preferred option over Vanilla VAE when it comes to exploring VAE latent space for the biological data analysis.

## Methods

### VAE: a general overview

Variational Autoencoder (VAE) is a deep generative model with a flavour of both neural networks and probabilistic graphical models[2]. It learns the latent representation (*z*) of the original space (data *x*). At first glance, we could regard the VAE model as a simple graphical model (Supplementary Figure S1). In the inference period, the latent variables (*z*) are sampled from a normal distribution, with fixed parameters (*θ*), new data *X* is generated; in the learning period, *z* and *X* are sampled N times and a neural network with a target (loss function) is trained to get the parameters *θ*, that is, the parameters of the neural networks. When we use VAE as an unsupervised learning method, only the learning period fits the purpose, unless we want to generate new data. After the training (the learning period), we get the latent representation of dataset (original space).

To further understand how VAE works without being distracted by the technical details, we can view VAE in this way (Figure 1). *P* (*X*) is the distribution of the original space, that is, the dataset; *q*_*ϕ*_(*z*| *x*) is the encoding distribution, via which we get *q*_*ϕ*_(*z*), the distribution of latent space; ideally, we sample *z* from a simple distribution *p*(*z*), for example, a normal distribution, then via a decoding distribution *p*_*θ*_(*x*| *z*), we get *P* (*X*′) (we call it the reconstruction space hereafter). *q*_*ϕ*_(*z*|*x*) and *p*_*θ*_(*x*|*z*) are coded by neural networks (*ϕ* and *θ* are the parameters of the encoder and decoder.). The training target includes two parts; the first part is “*q*_*ϕ*_(*z*) should be as similar as possible to *p*(*z*), the simple distribution, say, a normal distribution”((A) in Figure 1); the second is “the original space *P* (*X*) and reconstruction space *P* (*X*′) should be the same”((B) in Figure 1). The training target can be represented in equation (1), where *D* is any strict divergence, meaning that *D*(*q*‖*p*) *≥* 0 and *D*(*q‖p*) = 0 if and only if *q* = *p, λ* > 0 is a scaling coefficient.

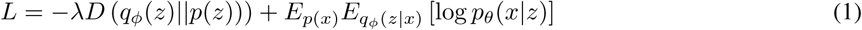

In the Vanilla VAE model, *D* is Kullback–Leibler divergence, and *λ* = 1 in the current study. so the first part of the formula 1 can be rewritten as *E*_*p*(*x*)_ [*-KL*(*q*_*ϕ*_(*z*|*x)‖p*(*z*))]

### MMD-VAE

Maximum Mean Discrepancy (MMD) [16][17] is defined to measure the discrepancy between two distributions; it works based on the presumption that two distributions are identical if and only if all moments are identical[18].

In practice, a biased empirical estimate of the MMD is used by calculating empirical expectations computed on the samples *X* and *Y*, by equation (2), where *F* is a class of functions *f* : *X*→ ℝ and *X* is the space where *x*_*i*_ and *y*_*i*_ are defined.

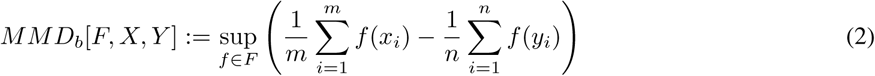

When it comes to the implementation, MMD can be calculated efficiently via the kernel embedding trick[18], as equation (3)

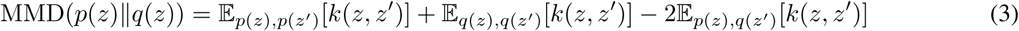

where *k*(*z, z*′) is any universal kernel, such as Gaussian 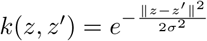.

The difference of MMD-VAE from the stand VAE model is the divergence in the training target. In MMD-VAE, MMD is used instead of KL divergence.

Intuitively speaking, in terms of the implementation of MMD-VAE, for the two distributions p(z) and q(z), as mentioned in equation (1) and (3), *MMD*(*p*(*z*), *q*(*z*)) is a value calculated representing this discrepancy. The smaller *MMD*(*p*(*z*), *q*(*z*)) is, the more similar *p*(*z*) and *q*(*z*) are.

All VAE architectures were implemented by TensorFlow [19] and TensorFlow Probability [20], visualised in TensorBoard, and run in Google Colaboratory (Colab) with GPU (Tesla K80) enabled.

### Datasets and data preprocessing

Single cell data usually mean single-cell RNA-seq data and mass cytometry (cyTOF) data. In this experiment, four published datasets were used; two are single-cell RNAseq data; two, mass cytometry data. These datasets are briefly summarized in Table 3. All of these datasets have been classified by the experts authoring the original papers using the computational methods described in these papers.

**Table 3:** An overview of the datasets used in the study. Four datasets were used: two single cell RNAseq data and two mass cytometry data.

| Dataset | Type | Cell No. | Class No. | Dimension | Supplement |
| --- | --- | --- | --- | --- | --- |
| SN | scRNA-Seq | 622 | 11 | 25334 | Sensory Neurons, 55 principal components were used |
| RBN | scRNA-Seq | 26830 | 18 | 13166 | Retinal Bipolar Neurons, 37 principal components were used |
| AML | Mass Cytometry | 104184 | 14 | 32 | AML benchmark dataset |
| PAN | Mass Cytometry | 102877 | 24 | 48 | Panorama benchmark dataset |

### Single-cell RNAseq datasets

Dataset SN is the single-cell transcriptome dataset of mouse neurons classified in the original paper [7] by sampling, unsupervised grouping and comprehensive transcriptome analysis; the data in use contain 622 cells, 11 classes and have 55 dimensions (principal components extracted from 25334 dimensions (genes))

Dataset RBN is the single-cell transcriptome dataset of mouse retinal bipolar cells classified in the original paper [23] using unsupervised clustering; the data in use contain 26830 cells, 18 classes and have 37 dimensions (principal components extracted from 13166 dimensions (genes)).

Both sing-cell RNAseq datasets have gone through Principal Component Analysis PCA (implemented in Python package: Scikit-learn [24]) first to reduce dimensions, and this has to be done because of the following three reasons: 1) Handling high-dimension-low-sample datasets has been a computational challenge for deep neural networks [25], which is not the main focus of this research and can be a hurdle for the experiment in this research 2) Even if it were not a challenge, the computation cost would still be significantly expensive, which Google Colab currently could not handle; 3) PCA has been a well-studied method for dimension reduction, and it is reasonable to assume that PCA does its job.

A permutation test [23] was reused to retain those principal components that capture statistically significantly correlated variation among the genes, which cannot be attributed to random “noise.” PCA was performed on *N* (*N* = 50 in current experiment) randomized versions of the data, in each version of which, all the columns of the original expression data frame (genes) were randomly and independently permuted; then a threshold eigenvalue is calculated (the average of the maximum eigenvalues in each version of the randomized data); in the original data, any principal component whose eigenvalue is greater than the threshold is kept.

PCA was implemented in a HPC cloud virtual machine (64GB memory 8 CPU) provided by SURFsara.

### Mass cytometry datasets

Dataset AML is a mass cytometry dataset of cryopreserved bone marrow aspirates from pediatric patients diagnosed with acute myeloid leukaemia. The data have been classified by the approach PhenoGraph from the original paper [21]. The data in use contain 104184 cells, 14 classes and have 32 dimensions.

Dataset PAN is a mass cytometry dataset from mice. The data have been classified (gated) by the algorithm “X-shift” from the original paper [22] already. The data in use contain 514386 cells, 24 classes and have 48 dimensions.

Both datasets have gone through the same preprocess where the data are rescaled by equation (4) where *N*_*original*_ is the orignal count of the dataset.

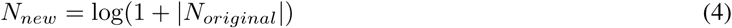

### Neural network classifier

To measure how much information has been retained in the latent space and reconstruction space of VAE, a neural network classifier is implemented in Keras to detect how many of the cells can still be correctly classified in the latent space and reconstruction space. Two metrics are used to measure the accuracy of the classifier: F2 score (F-measure in equation (5)) and Matthews correlation coefficient (equation (6)).

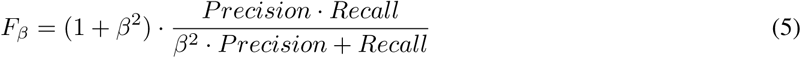

where *β* = 2 and 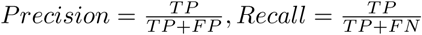.

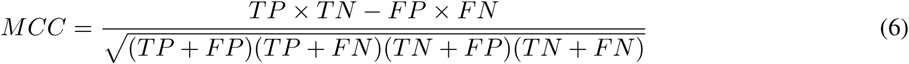

TP = True Positive; FP = False Positive; FN = False Negative; TN = True Negative

A permutation test is also conducted to show that a neural network classifier only works over the original data, not the permuted data. For each permuted version of the dataset, the classification accuracy in terms of the two metrics mentioned above is on the random level. So the classifier can be confidently used.

The code is implemented by Keras 2.2.4[26] with Tensorflow[19] as the backend and run in Google Colab with GPU (Tesla K80) enabled.

### Experiment strategy

The parameter optimization can add much uncertainty to experiment to compare the two VAE architectures, therefore a proper strategy is needed to exercise the control over the uncertainty. In this study, VAE and neural network classifier go hand in hand in the mental model (see the discussion). Parameters has to be optimized dataset by dataset. For each dataset, parameters need training only on the side of Vanilla VAE (including Vanilla VAE and its corresponding three neural classifiers) (Figure 2); afterwards, the parameters are applied to MMD-VAE and its corresponding three neural network classifiers over the same dataset. The training was done manually. The intuition of determining the neural network archecture (the number of layers, etc.) was referred to the publications[**?**] [**?**]. Dimension of latent space was tried in the range of 3,4,5,10; learning rate was tried in the range of 2e-2, 1e-2, 1e-3, 1e-4, 1e-5; batch size was tried in the range of 128 and 256. A complete list of the optimized parameters can be found in Supplementary Table S2 (VAE) and S3 (Classifier).

After the parameter optimization, Each VAE + neural network classifier is run five times over each dataset; due to the stochastic nature of VAE, it occasionally delivers bad results which can be detected intuitively by humans; this unlucky run is excluded from the final result.

The neural network classifier is trained on 22.5% of the dataset; 2.5% is used as the validation dataset; and 75% as the test dataset.

## Data availability

Data and code that contribute to the reproducibility of the results in this manuscript, as well as other technical details, are available at: https://research-project.gitlab.io/mmd-vae

## Acknowledgements

I thank Peter Bloem for giving feedback on this manuscript. I thank Hui Liu for proofreading. I thank Jaap Heringa and Sanne Abeln for the support during the research period.

## Author contributions statement

C.Z. is the only author who contributes to every aspect of this work.

