## Supplementary tables and figures for "Single-Cell Data Analysis Using MMD Variational Autoencoder for a More Informative Latent Representation"

### SUPPLEMENTARY INFORMATION

A PREPRINT

**Chao Zhang***
Department of Computer Science
Vrije Universiteit Amsterdam
Amsterdam, 1081 HV


June 23, 2019

**ABSTRACT**

Variational Autoencoder (VAE) is a generative model from the computer vision community; it learns a latent representation of images and generates new images in an unsupervised way. Recently, Vanilla VAE has been applied to single-cell data analysis, in the hope of harnessing the representation power of latent space to evade the "curse of dimensionality" of the original dataset. However, Vanilla VAE is suffering from the issue of less informative latent space, which raises a question concerning the reliability of Vanilla VAE latent space in representing the high-dimensional single-cell datasets. Therefore I set up such a study to examine this issue from the multiple perspectives.

This paper confirms the issue of Vanilla VAE by comparing it with MMD-VAE, a variant of VAE which has claimed to have overcome this issue based on image data, across a series of single-cell RNAseq and mass cytometry datasets. The result indicates that MMD-VAE is superior to Vanilla VAE in retaining the information not only in the latent space but also the reconstruction space, which suggests that MMD-VAE be a better option for single-cell data analysis than Vanilla VAE.

***Keywords*** variational autoencoder · single-cell RNA seq · mass cytometry · deep learning · artificial intelligence · bioinformatics

#### Supplementary Table

##### Supplementary table S1

| Dataset | F2-Score | MCC |
|---|---|---|
| AML | 0.970 | 0.972 |
| PAN | 0.990 | 0.991 |
| SN | 0.794 | 0.793 |
| RBN | 0.982 | 0.983 |

Table 1: The accuracy of the classifier applied to the original datasets, two metrics are used: F2-Score and MCC.

*https://hi-it.org

**Supplementary Table S2**

| vanilla VAE/MMD-VAE | | | | | | |
|---|---|---|---|---|---|---|
| Dataset | dimension of latent space | training steps | en/decoder structure | learning rate | Activation Function | batch size |
| AML | 3 | 10,000 | [24, 16, 8], [8, 16, 24] | 1e-3 | Relu | 256 |
| PAN | 3 | 12,000 | [36, 18, 9], [9, 18, 36] | | | |
| SN | 10 | 10,000 | [40, 24, 8], [8, 24, 40] | | | |
| RBN | 3 | 10,000 | [24, 12, 6], [6, 12, 24] | | | |

Table 2: The parameters of variational autoencoder; parameters were manually optimized over Vanilla VAE per dataset (together with the neural network classifier) and re-applied to MMD-VAE accordingly.

**Supplementary Table S3**

| Neural Network Classifier | | | | | | | | |
|---|---|---|---|---|---|---|---|---|
| Dataset | VAE Space | structure | dropout | epoch | learning rate | l2 regularization | Activation Function | batch size |
| AML | Recons/Latent | [12,6] | 0 | 20 | 0.001 | 0 | Relu | 128 |
| PAN | Reconstruction | [36,18,9] | 0 | 20 | 0.001 | 0.01 | | |
| | Latent | [36,24,12,6] | | | 0.1 | 0 | | |
| SN | Reconstruction | [40,15] | 0.1 | 500 | 0.02 | 0.01 | | |
| | Latent | [12,6,3] | 0 | 1000 | | 0 | | |
| RBN | Reconstruction | [20,10,5] | 0 | 100 | 0.001 | 0 | | |
| | Latent | [12,6,3] | | 200 | 0.02 | | | |

Table 3: The parameters of neural network classifier for the original space, latent space and reconstruction space per dataset, parameters were manually optimized over the output of Vanilla VAE per space per dataset and re-applied to each space of MMD-VAE accordingly.

#### Supplement figure

##### Supplementary Figure S1

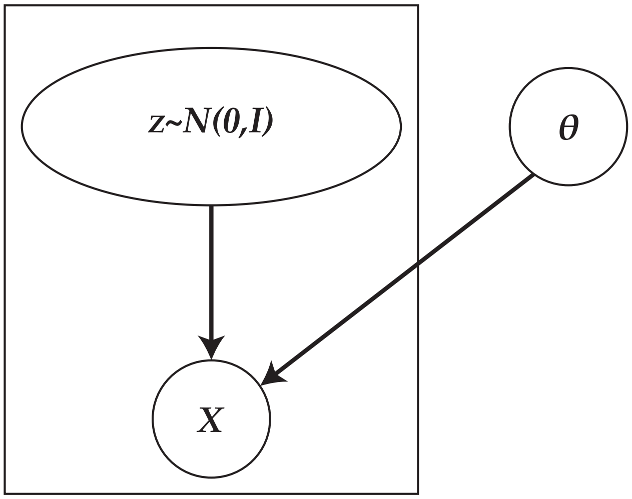


Figure 1: A graphical model representing Variational Autoencoder (VAE): in inference period, $z\ N(0, I)$ represents the latent space characterized by a normal distribution, $X$ represents the construction space generated by the sampled $z$ and the neural network architecture; $\theta$ represents the fixed parameters confined by neural network architecture.

#### Data availability

Data and code that contributes to the reproducibility of the results in this manuscript, as well as other technical details, are available at: https://research-project.gitlab.io/mmd-vae/
